# Rhythmic Cilium in SCN Neuron is a Gatekeeper for the Intrinsic Circadian Clock

**DOI:** 10.1101/2022.01.26.477948

**Authors:** Hai-Qing Tu, Sen Li, Yu-Ling Xu, Yu-Cheng Zhang, Xiao-Xiao Jian, Guang-Ping Song, Min Wu, Zeng-Qing Song, Huai-Bin Hu, Pei-Yao Li, Li-Yun Liang, Jin-Feng Yuan, Xiao-Lin Shen, Jia-Ning Li, Qiu-Ying Han, Kai Wang, Tao Zhang, Tao Zhou, Ai-Ling Li, Xue-Min Zhang, Hui-Yan Li

## Abstract

The internal circadian rhythm is controlled by the central pacemaker in the hypothalamic suprachiasmatic nucleus (SCN). SCN drives coherent and synchronized circadian oscillations via intercellular coupling, which are resistant to environmental perturbations. Here we report that primary cilium is a critical device for intercellular coupling among SCN neurons and acts as a gatekeeper to maintain the internal clock in mice. A subset of SCN neurons, namely neuromedin S-producing (NMS) neurons, exhibit cilia dynamics with a pronounced circadian rhythmicity. Genetic ablation of ciliogenesis in NMS neurons enables a rapid phase shift of the internal clock under experimental jet lag conditions. The circadian rhythms of individual neurons in cilia-deficient SCN slices lose their coherence following external perturbations. Rhythmic cilia dynamics drive oscillations of Sonic Hedgehog (Shh) signaling and oscillated expressions of multiple circadian genes in SCN neurons. Genetic and chemical inactivation of Shh signaling in NMS neurons phenocopies the effect of cilia ablation. Our findings establish ciliary signaling as a novel interneuronal coupling mechanism in the SCN and may lead to novel therapy of circadian disruption-linked diseases.

**One-Sentence Summary:** Rhythmic cilium is a critical device for intercellular coupling among SCN neurons and acts as gatekeeper for the internal clock.

## Introduction

All mammals possess an internal circadian clock (about 24 hours) that regulates daily oscillations in metabolism, physiology and behavior, such as rest-activity and sleep-wake cycles (*1*). The suprachiasmatic nucleus (SCN) acts as the master pacemaker of the internal circadian clock (*2, 3*). Its autonomous and coherent oscillatory output signals orchestrate the peripheral clocks in multiple tissues throughout the body (*4, 5*). Environmental circadian disruptions, such as acute jet lag and long-term shift work, cause temporal unsynchronization between the internal circadian clock and external time cues, leading to physiological stress (*6*). Circadian disruption has been implicated in tumorigenesis and various psychiatric, neurological and metabolic diseases, including depression and diabetes (*7, 8*).

The SCN contains a heterogenous population of approximately 20,000 neurons, most of which can individually generate autonomous circadian oscillations (*9, 10*). These intracellular processes are driven by autoregulatory transcription–translation feedback loops (TTFLs) of clock genes (*11, 12*). The function of SCN as the master pacemaker relies on intercellular coupling, a process that synchronizes period and phase among SCN neurons (*13*). Intercellular coupling is unique to SCN neurons and enables the SCN to generate robust and coherent oscillations at the population level, which are resistant to environmental perturbations (*14, 15*). Several neurotransmitters, including vasoactive intestinal peptide (VIP), gamma-aminobutyric acid (GABA) and arginine vasopression (AVP), play key roles in maintaining this intercellular coupling (*16-20*). The release of these neurotransmitters is regulated by clock genes, whose transcription can be further activated by these neurotransmitters (*21*). Thus, these neurotransmitters and clock genes form a feed forward loop to maintain intercellular coupling in the SCN.

The primary cilium, a sensory organelle nucleated by the mother centriole, plays a crucial role in mammalian embryonic development (*22*). Defective ciliogenesis results in a series of human disorders collectively known as ciliopathies (*23, 24*). The primary cilium is also present in adult neurons, and involved in the regulation of glycometabolism (*25*). Here we find that a subset of SCN neurons exhibit cilia dynamics with a pronounced circadian rhythmicity, and further identify primary cilia as a critical device for intercellular coupling to maintain the circadian clock in the SCN.

## Results

We examined the distribution of primary cilia in the mouse brain, and found that many SCN neurons contained primary cilia (Fig. 1, A and B and movie S1). Because the SCN is the master pacemaker of circadian clock, we tested whether primary cilia of SCN neurons exhibited oscillatory rhythms during light-dark (LD) and dark-dark (DD) cycles. When mice were maintained in LD cycles, both the number and length of primary cilia showed a pronounced circadian rhythmicity, peaking at ZT 0 and reaching trough at ZT 12 (Fig. 1, C to E) (ZT represents Zeitgeber time used in LD cycles; ZT 0 and ZT 12 are lights on and off, respectively). This oscillation was antiphase to that of clock gene *Cry1*. Rhythmic oscillations of primary cilia in the SCN were also observed in DD cycles, peaking at CT 0 and reaching trough at CT 12 (Fig. 1F, and fig. S1, A and B) (CT represents circadian time; CT 0 and CT 12 are subjective dawn and dusk, respectively), indicating that these ciliary oscillations are not driven by light but rather internal rhythms. We next used a transgenic mouse model expressing ADP ribosylation factor like GTPase 13B (ARL13B)-mCherry fusion protein to label the axoneme of primary cilia, and also observed the diurnal oscillations of primary cilia in the SCN during DD cycles (fig. S1C). Strikingly, the primary cilia in other cerebral regions and peripheral tissues including the paraventricular nucleus of the hypothalamus (PVN), hippocampus, kidney and pancreas, lack circadian rhythmicity (Fig. 1G, and fig. S1, D to G). Bmal1 is a core component of the mammalian circadian clock, and its deletion expectedly abolished circadian behaviors in mice (*26, 27*) (fig. S1H). We found that the circadian oscillation of cilia was lost in Bmal1-deficient mice, indicating that the rhythmicity of cilia is regulated by clock genes (Fig. 1H).

**Fig. 1.**
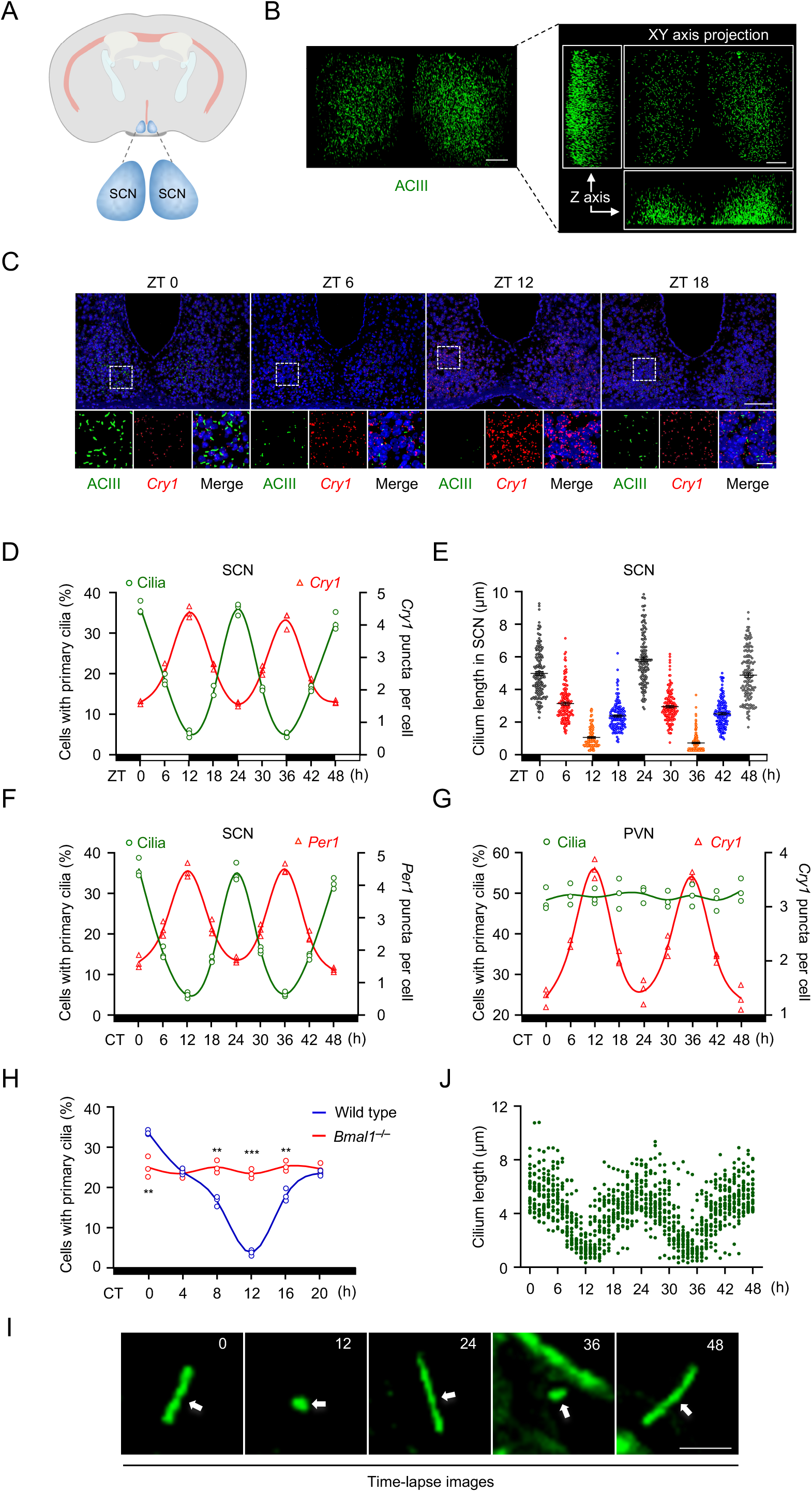
Primary cilia in the SCN exhibit circadian dynamics. (**A**) Schematic diagram of the suprachiasmatic nucleus (SCN). (**B**) Representative three-dimensional reconstructed projection images of the SCN at 20× magnification of the two-photon imaging are shown. SCN slices were stained with anti-ACIII antibody (primary cilia marker, green). Scale bars, 50 μm. (**C**) Representative images of primary cilia and expression of clock gene *Cry1* in the SCN during light-dark (LD) cycle. SCN coronal sections were stained with anti-ACIII antibody (green), *Cry1* RNAscope probes (red) and Hoechst (blue). Insets show zoomed-in views of the boxed regions in the core of the SCN. Scale bars, 100 μm (main image) and 20 μm (magnified region). (**D**) The percentage of cells with primary cilia or *Cry1* RNAscope signals in the SCN were determined based on (C) (*n* = 3 mice per time point). (**E**) Quantitative analysis of the cilium length in (C). Each dot represents one cell. Data are presented as mean ± SEM. (**F**) The percentage of cells with primary cilia or *Per1* RNAscope signals in the SCN were quantified during constant darkness (DD) cycle (*n* = 3 mice per time point). (**G**) The percentage of cells with primary cilia or *Cry1* RNAscope signals in the PVN were determined during DD cycle (*n* = 3 mice per time point). PVN, paraventricular nucleus. (**H**) The percentage of cells with primary cilia in the SCN for wild type and *Bmal1*^**–/–**^ mice in DD cycle (*n* = 3 mice per time point). Statistics indicate significance by unpaired *t*-test (I). \*\**P* < 0.01, \*\*\**P* < 0.001. (**I**) Representative time-lapse images of the primary cilium in SCN neurons for 48 hours. Isolated live SCN neurons from postnatal mice were transduced with modified baculovirus encoding mCherry tagged, constitutively ciliary-localized protein, 5-hydroxytryptamine receptor 6 (5-HT6). The numbers on the images indicate the time in hours. Arrows indicate primary cilia. Scale bar, 5 μm. (**J**) Quantitative analysis of the cilium length in (I). Each dot represents one cell.

To monitor the dynamics of primary cilia, we isolated live SCN neurons from postnatal mice and transduced them with modified baculovirus encoding mCherry tagged, constitutively ciliary-localized protein, 5-hydroxytryptamine receptor 6 (5-HT6), and performed time-lapse imaging. Consistent with fixed-slice data, primary cilia indeed display circadian rhythmic oscillation: they took approximately 12 hours to shorten to the minimum length, and regrew to the maximum length during the next 12 hours (movie S2 and Fig. 1, I and J, and fig. S1I). We also observed the low expression of Cry1 in ciliated cells as compared to non-ciliated cells (fig. S1J), consistent with our *in vivo* observations.

The SCN consists of multiple types of neurons, including AVP-expressing neurons, VIP-expressing neurons and NMS-expressing neurons. NMS neurons represent 40% of all SCN neurons and encompass the majority of VIP- and AVP-expressing neurons (*28*). To identify the cell types of ciliated neurons in the SCN, we crossed *Nms*-*Cre* or *Vip*-*Cre* mice with *Rosa-stop-tdTomato* reporter mice to label NMS or VIP neurons. As revealed by ACIII staining of SCN coronal sections, we found that 90% of ciliated neurons were NMS neurons, and 31% were VIP neurons (Fig. 2, A and B). We then performed double immunostaining using anti-ACIII and anti-AVP antibodies and found that only 5% of ciliated cells were AVP neurons (Fig. 2, A and B). Thus, primary cilia are mainly present on NMS neurons in the SCN (Fig. 2C).

**Fig. 2.**
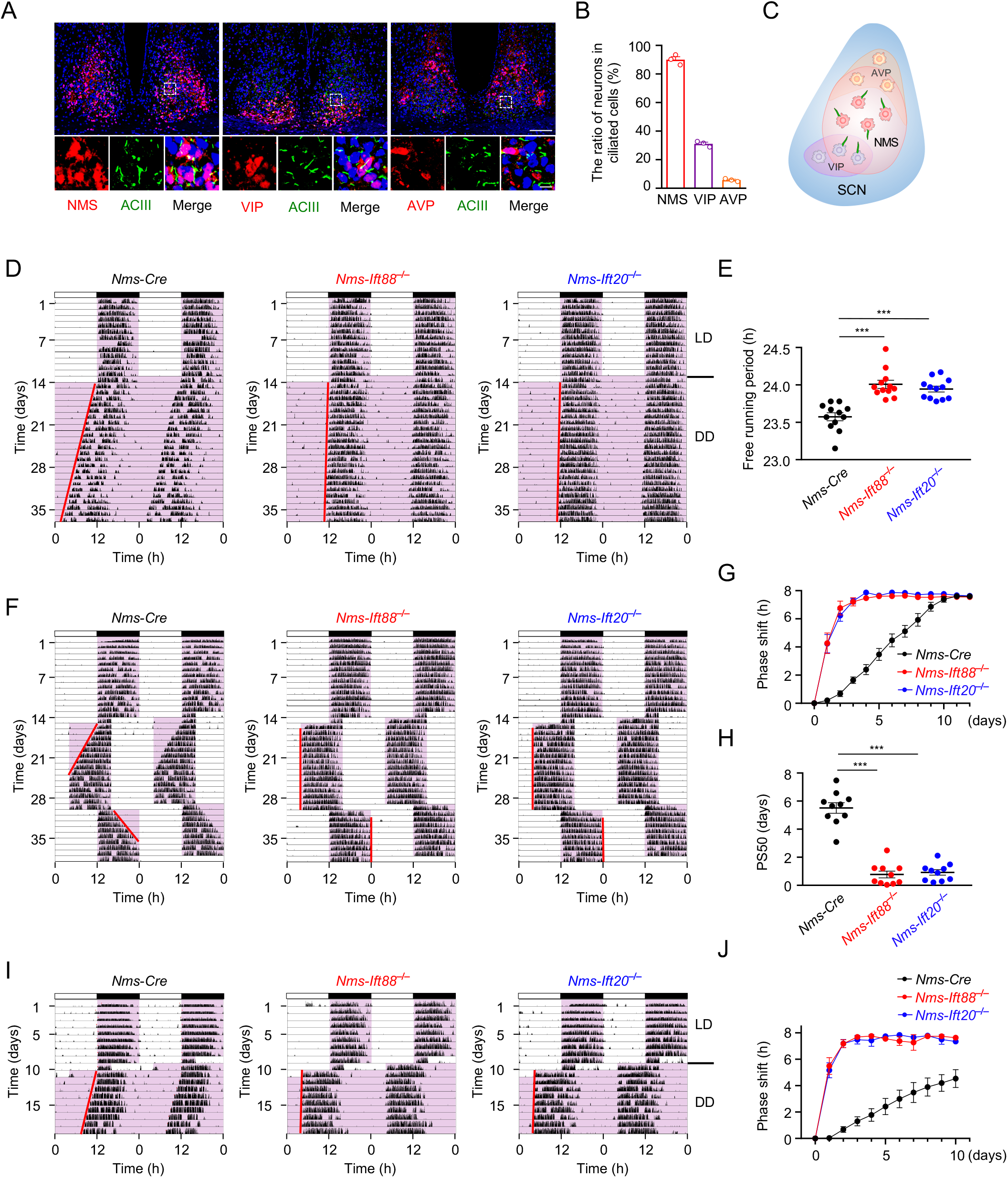
Primary cilia in NMS neurons confer robustness to the intrinsic circadian clock. (**A**) Representative images of primary cilia in multiple types of SCN neurons. For NMS and VIP neurons, SCN coronal sections from *Nms*-*tdTomato* or *Vip*-*tdTomato* mice were stained with anti-ACIII antibody (green) and Hoechst (blue). For AVP neurons, SCN coronal sections were stained with anti-ACIII antibody (green), anti-AVP antibody (red) and Hoechst (blue). Insets show zoomed-in views of the boxed regions in the core of the SCN. Scale bars, 100 μm (main image) and 10 μm (magnified region). (**B**) Quantitative analysis of the ratio of NMS, VIP or AVP neurons in total ciliated SCN cells in (A) (*n* = 3). (**C**) The distribution diagram of primary cilia in NMS, AVP and VIP neurons. (**D**) Representative double-plotted actograms of wheel-running activities in *Nms-Cre* and SCN^cilia-null^ (*Nms-Ift88*^−***/***−^ and *Nms-Ift20*^−***/***−^) mice. Activity records are double plotted so that 48 h are represented horizontally. Each 24 h interval is both presented to the right of and beneath the preceding day. Animals were initially housed in LD conditions for 14 days and were then transferred to DD cycle. (**E**) Quantification of free wheel-running period in (D) (*n* = 12). (**F**)Representative double-plotted actogram of *Nms-Cre* and SCN^cilia-null^ mice under experimental jet-lag conditions. Mice were maintained in LD cycles for 14 days, then the cycle was advanced 8 h, and 15 days later, the cycle was returned to the original lighting regime. (**G**) Line graphs showing the daily phase shift of wheel-running activities after an 8 h advance in (F) (*n* = 10). (**H**) PS50 (50% phase-shift) values after an 8 h advance in (F) (*n* = 10). (**I**) Representative double-plotted actograms of *Nms-Cre* and SCN^cilia-null^ mice subjected to an 8 h phase advance on day 1 and released to DD. (**J**) Line graphs showing the daily phase shift of wheel-running activities in (I)(*n* = 10). For this and subsequent figures, wheeling-running activity is indicated by black markings. White and pink background indicates lights on and off, respectively. The red lines on the actograms delineate the phase of activity onset or offset. All data are presented as mean ± SEM. Statistics indicate significance by Kruskal-Wallis test (E), one-way ANOVA with Dunnett correction (H). \*\*\**P* < 0.001.

We disrupted primary cilia specifically in NMS neurons by conditionally deleting *Ift88* or *Ift20*, two genes required for ciliogenesis (*29, 30*). Primary cilia on SCN neurons were abolished in *Nms-Ift88*^−***/***−^ or *Nms-Ift20*^−***/***−^ mice, referred to hereafter as SCN^cilia-null^ mice (fig. S2, A to D). In contrast, the number of primary cilia in other tissues were comparable between control and SCN^cilia-null^ mice (fig. S2, E and F). Deletion of *Ift88* or *Ift20* in NMS neurons did not affect mouse development and the morphology of the SCN (fig. S3, A to E). Next, we examined whether primary cilia contribute to the pacemaker function of SCN by monitoring locomotor activity of SCN^cilia-null^ mice. Under LD cycles, both control and SCN^cilia-null^ mice exhibited normal locomotor activity with a wheel-running period of 24 h (Fig. 2D). Under DD cycles, control mice exhibited intrinsic periods of 23.6 ± 0.1 h (Fig. 2, D and E). SCN^cilia-null^ mice had a moderatelly elongated intrinsic period (24.0 ± 0.1 h for *Nms-Ift88*^−***/***−^ mice and 23.9 ± 0.1 h for *Nms-Ift20*^−***/***−^ mice) (Fig. 2, D and E).

We further examined the behavior of SCN^cilia-null^ mice under experimental jet-lag conditions. Mice were maintained in normal LD cycles for 14 days; then the cycle was advanced by 8 h; and 15 days later, the cycle was returned to the original lighting regime. Under LD cycles, both control and SCN^cilia-null^ mice successfully entrained to the LD schedule. When the lighting cycle was advanced, control mice re-entrained progressively over 9–11 days (Fig. 2, F and G). In contrast, SCN^cilia-null^ mice re-entrained more quickly, and the entrainment was complete within 1–3 days (Fig. 2, F and G). The 50% phase-shift value (PS_50_) of the control mice was 5.5 ± 0.4 days, whereas the PS_50_ values of *Nms-Ift88*^−***/***−^ and *Nms-Ift20*^−***/***−^ mice were 0.7 ± 0.2 and 0.9 ± 0.2 days, respectively (Fig. 2H). When the cycle was returned to the original lighting regime, control mice re-entrained progressively over several days (Fig. 2F, fig. S3, F and G), whereas SCN^cilia-null^ mice again re-entrained immediately to the LD cycle. There were no significant differences between control and SCN^cilia-null^ mice in the number of *c-Fos*-positive cells induced by a 30-min light pulse at CT 22 (fig. S3, H and I), indicating the light response of SCN^cilia-null^ mice was similar to that of control mice. These results demonstrate that SCN^cilia-null^ mice are more adaptive to phase shifts during LD cycles, suggesting that primary cilia confer resistance to environmental time cues.

To exclude the effect of light on locomotor activity of mice, we subjected mice to constant darkness (DD) after an 8-hour phase advance following LD cycles. Under DD, SCN^cilia-null^ mice exhibited significant phase advances in the free-running behavior, whereas control mice did not (Fig. 2, I and J). This finding strongly suggests that the immediate adaptation to a new LD cycle in SCN^cilia-null^ mice is a rapid phase shift of the internal clock, not due to a masking effect of the environmental LD cycle.

SCN neurons rely on intercellular coupling to maintain intrinsic circadian behavior. Intercellular coupling confers robustness to neuronal networks and synchronizes periods of individual cellular oscillators. *Per2::Luciferase* (*Per2::Luc*) transgenic reporter mice can be used to track Per2 rhythmic expression in single cells *ex vivo*. To test whether primary cilia are required for intercellular coupling among SCN neurons, we performed real-time luciferase luminescence imaging of SCN slices isolated from control or *Nms-Ift88*^−***/***−^ mice that expresssed the Per2::Luc reporter. As expected, the circadian rhythmicity of the analyzed SCN cell population was coherent in both control and *Nms-Ift88*^−***/***−^ slices under normal cultured conditions (Fig. 3, A to D). We then applied tetrodotoxin (TTX), a sodium ion channel blocker, to disrupt the intercellular coupling of SCN neurons. TTX disrupted the phase order of both control and *Nms-Ift88*^−***/***−^ SCN neurons (Fig. 3, A to D). After the washout of TTX, control neurons recovered their coherent phase order, whereas *Nms-Ift88*^−***/***−^ neurons failed to do so (Fig. 3, A to D, and fig. S4, and movie S3 and S4). This result suggests that primary cilia in NMS neurons promote intercellular coupling.

**Fig. 3.**
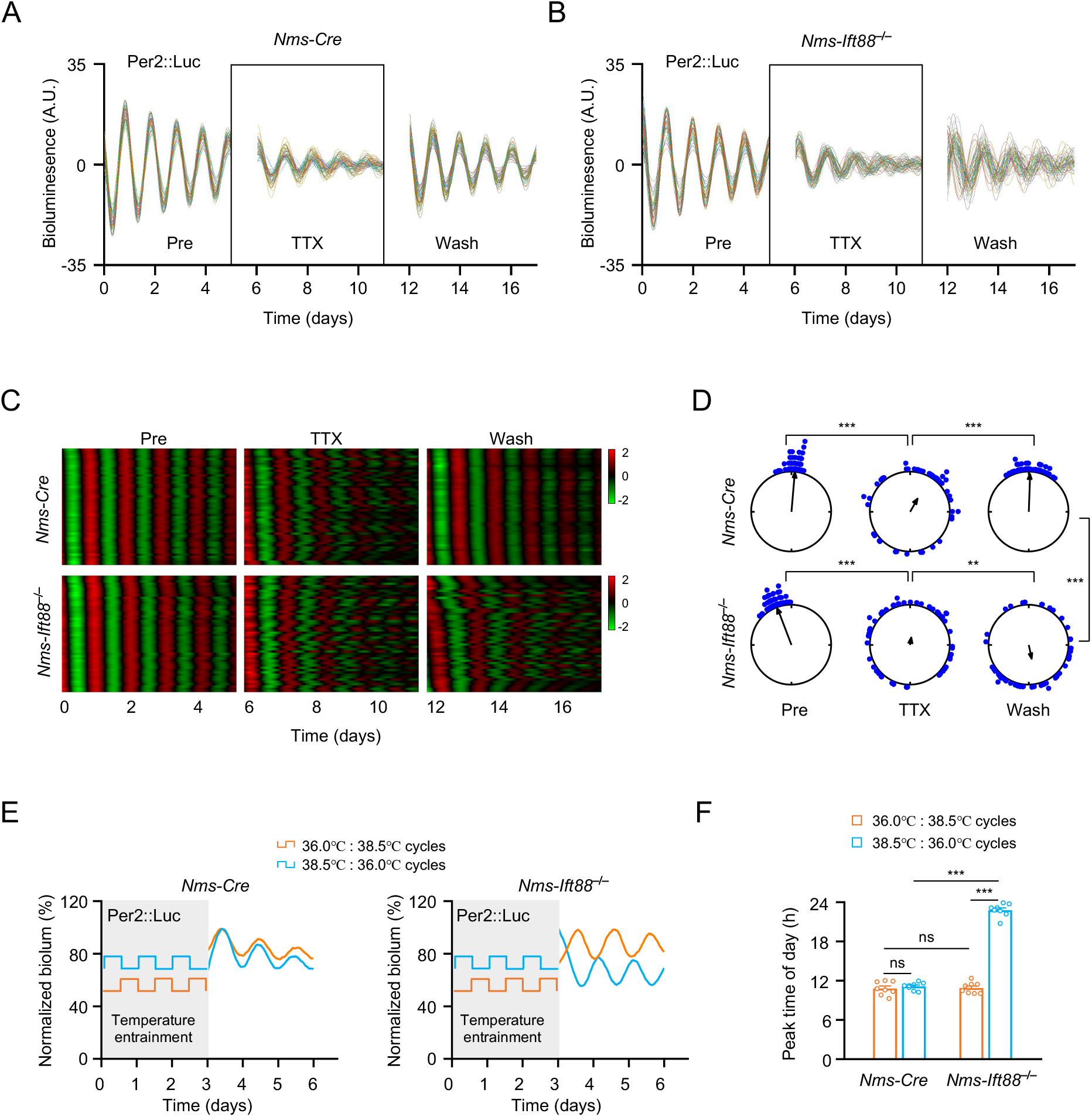
Primary cilia promote interneuronal coupling in the SCN. (**A** and **B**) Representative records of single-cell Per2 bioluminescence in *Nms-Cre* (A) and *Nms-Ift88*^−***/***−^ SCN slices (B) under three conditions: pre-treatment, TTX treatment, washout of TTX (*n* = 50 cells). (**C**) Representative heatmap of Per2 bioluminescence oscillation in (A) and (B). The data were ordered by phases, with earlier phases placed at the top. The red and green color represent high and low bioluminescence intensity, respectively. (**D**) Rayleigh plots show the phase distribution of single cells during the third day (midpoint) in (C). The scattering of dots represents the phase distributions of single neurons. Arrows represent mean circular phase, and the length of the arrow represents the strength of synchronization. (**E**) Representative records of Per2 bioluminescence rhythms in *Nms-Cre* and *Nms-Ift88*^−***/***−^ SCN slices. SCN slices were exposed to 12 h of 36°C and 12 h of 38.5°C temperature cycles or oppositely phased temperature cycles for three days, and their bioluminescence was then monitored continuously at a constant 36°C. Bioluminescence was normalized to the first peak. (**F**) Quantitative analysis of the peak time of Per2 bioluminescence after the temperature cycle (*n* = 8 for each group). Data are presented as mean ± SEM. Statistics indicate significance by Watson-Wheeler test (D), two-way ANOVA with Bonferroni correction (F). \*\**P* < 0.01, \*\*\**P* < 0.001, ns, not significant.

Intercellular coupling in the SCN is required for the resistance to physiological temperature changes (*15*). We then tested whether primary cilia in the SCN contributed to the resistance to cyclic temperature entrainment. Control and *Nms-Ift88*^−***/***−^ SCN slices were exposed to 12 h of 36°C and 12 h of 38.5°C temperature cycles (normal body temperature cycles) or oppositely phased temperature cycles for three days. Both control and *Nms-Ift88*^−***/***−^ SCN slices maintained their phases of Per2 bioluminescence after normal cyclic temperature entrainment. After oppositely phased temperature cycles, the phase of control SCN slices still remained unchanged, but the phase of *Nms-Ift88*^−***/***−^ SCN slices was obviously shifted (Fig. 3, E and F). This finding suggests that cilia-null SCN was less resistant to cyclic temperature changes. This resistance might be due to cilia-dependent intercellular coupling in the SCN.

The primary cilium is a critical organelle that regulates Sonic Hedgehog (Shh) signaling (*31-33*). In a whole-genome screen, inhibition of Hedgehog pathway resulted in long period length of circadian oscillations in U2OS cells (*34*). We also revealed that *Shh, Ptch1, Gli1* mRNA were indeed expressed in ciliated SCN neurons (fig. S5A). Furthermore, we found that the expression of Gli1 and Ptch1, two Shh signaling target genes, in the SCN exhibited rhythmic oscillation, which was lost in *Nms-Ift88*^−***/***−^ mice (fig. S5B). Thus, we next investigated whether Shh signaling was involved in the regulation of molecular rhythms and locomotor behaviors. The enrichment of smoothened (SMO) on cilia is required to initiate transduction of Hedgehog pathway (*35*). We treated Per2::Luc SCN slices with smoothened (SMO) inhibitor Vismodegib. The data shown that Vismodegib strongly suppressed circadian Per2::luciferase activity in SCN slices, and elicited a reversible dose-dependent reduction in the amplitude and lengthening in the period of Per2::Luc oscillations (fig. S6, A to C).

Next, we generated *Nms*-*Smo*^−***/***−^ mice to block Shh signaling in the SCN (fig. S7A). Deletion of Smo in NMS neurons did not affect mouse development (fig. S7, B to E). We then monitored the circadian behavior of *Nms*-*Smo*^−***/***−^ mice under experimental jet-lag conditions. *Nms*-*Smo*^−***/***−^ mice re-entrained immediately to the LD cycle, whereas control mice re-entrained progressively over several days (Fig. 4A and fig. S7, F and G). We further applied Vismodegib to the SCN through an osmotic minipump in live animals during experimental jet lag conditions. Vismodegib also caused immediate re-entrainment to the LD cycle (Fig. 4, B and C, and fig. S7H). These results demonstrate that Shh signaling is required for the resistance to environmental time cues.

**Fig. 4.**
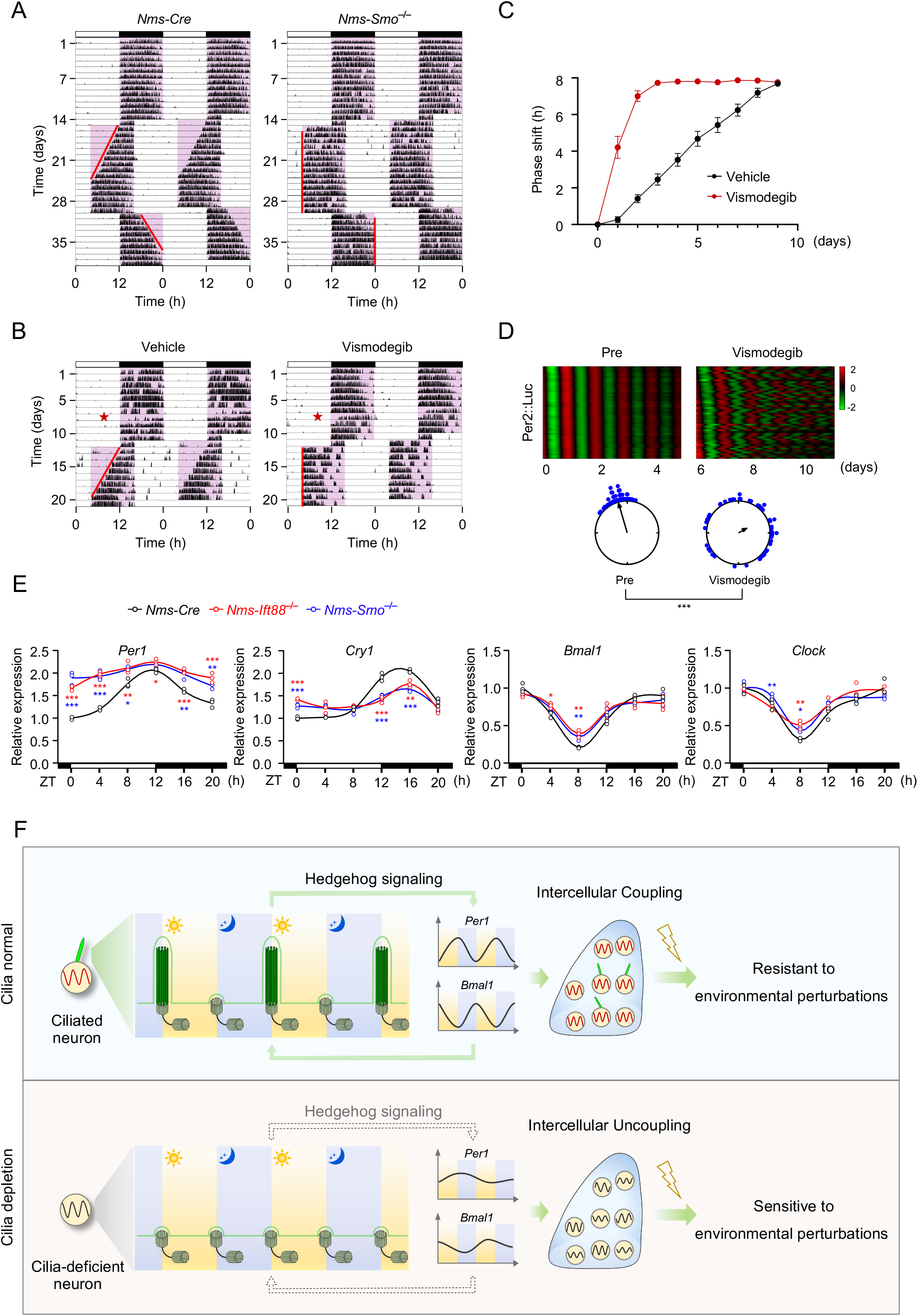
Hedgehog signaling maintains circadian rhythm in the SCN. (**A**) Representative double-plotted actogram of *Nms-Cre* and *Nms-Smo*^−***/***−^ mice under experimental jet-lag conditions. (**B**) Representative double-plotted actograms of mice treated with vehicle (left) or 5 mM Vismodegib (right) under experimental jet-lag conditions. Vismodegib was applied to the SCN by osmotic minipump. Asterisks indicate time of surgery. Three days after surgery, LD cycles were advanced by 8 h. The red lines on the actograms delineate the phase of activity onset. (**C**) Line graphs showing the daily phase shift of wheel-running activities after an 8 h advance in (B) (*n* = 11 for vehicle group, *n* = 12 for Vismodegib group). (**D**) Heatmap of Per2 bioluminescence oscillation and Rayleigh plots of phase distribution of single cells during the third day (midpoint) under pre- and post-treatment with 10 μM Vismodegib (*n* = 50 cells). (**E**) Quantitative real-time PCR analysis of core clock genes in the SCN from *Nms-Cre, Nms-Ift88*^−***/***−^ or *Nms-Smo*^−***/***−^ mice. SCN were collected at 4 h intervals across the LD cycle (*n* = 3 independent experiments). (**F**) Summary diagram showing primary cilia in SCN neurons and downstream Hedgehog signaling as critical regulatory mechanisms that promote interneuronal coupling, thereby maintaining SCN network synchrony and circadian rhythms. All data are presented as mean ± SEM. Statistics indicate significance by Watson-Wheeler test (D), one-way ANOVA with Dunnett correction (E). \**P* < 0.05, \*\**P* < 0.01, \*\*\**P* < 0.001.

To test whether Shh signaling plays a role in interneuronal coupling in the SCN, we analyzed circadian rhythms in SCN slices treated with Vismodegib. Single-cell bioluminescence imaging showed that Vismodegib disrupted the phase order of Per2::Luc (Fig. 4D, and fig. S8), suggesting that Shh signaling promotes intercellular coupling among SCN neurons. Since circadian clocks control the transcriptional expression of multiple factors important for interneuronal communications, we next performed qPCR with the isolated SCN from control, *Nms*-*Ift88*^−***/***−^ and *Nms*-*Smo*^−***/***−^ mice sacrificed every 4 h during the circadian cycle. The rhythmicity of core clock genes including *Per1, Cry1, Bmal1* and *Clock*, was altered in *Nms*-*Ift88*^−***/***−^ and *Nms*-*Smo*^−***/***−^ mice (Fig. 4E). These data suggest that primary cilia regulate circadian genes through the Shh pathway.

The SCN drives coherent and synchronized circadian oscillations that are resistant to environmental perturbations, and this resistance relies on interneuronal coupling. Our findings establish primary cilia in NMS neurons and downstream Shh signaling as critical regulatory mechanisms that promote interneuronal coupling, thereby maintaining SCN network synchrony and circadian rhythms (Fig. 4F). The rhythmic cilia dynamics in the SCN lead to rhythmic oscillation of Shh signaling, which in turn drives rhythmic expression of core clock genes. This feed forward loop sustains robust and coherent oscillation at the cell population level, making the intrinsic clock resistant to environment perturbations. In the future, it will be interesting to determine how clock genes drive cilia dynamics in the SCN and how Shh signaling regulates the expression of clock genes.

Epidemiological studies have linked frequent cross-time zone travel and shift work to high blood pressure, obesity and other metabolic disorders. Our results show that pharmacological blockade of the Shh pathway accelerates recovery from experimental jet lag in mice. Targeting Shh signaling might be a potential therapeutic strategy for the treatment of human diseases related to circadian disruptions.

## Acknowledgments

We thank Eric Erquan Zhang for the gift of *Bmal1*^−***/***−^ mice and *Per2::Luciferase* transgenic reporter mice (Jackson Laboratory). We thank Minmin Luo for technique support of intracerebroventricular injections. We thank the Center of Biomedical Analysis, Tsinghua University for the two-photon microscopy imaging.

## Author contributions

H.Y.L. and X.M.Z. supervised the project; H.Q.T.,S.L. and Y.L.X. built the experimental system and carried out most of the experiments; Y.C.Z. conducted locomotor activity assays; X.X.J. and G.P.S. performed the experiments of animal genotyping; M.W., Z.Q.S., H.B.H., P.Y.L. and L.Y.L. analyzed the data and amended the original draft of the manuscript; J.F.Y., X.L.S., J.N.L., Q.Y.H., K.W. and T.Z. provided reagents and suggestions; T.Z. and A.L.L. carried out the statistics; H.Q.T., Y.L.X., S.L., H.Y.L. and X.M.Z. wrote the paper. All authors discussed the results and commented on the manuscript.

## Data and materials availability

All data are available in the manuscript or the supplementary materials.

## Supplementary Materials

Materials and Methods

Figs. S1 to S8

Tables S1 to S2

References (*36–40*)

Movies S1 to S4

