## Supplementary Materials for "Rhythmic Cilium in SCN Neuron is a Gatekeeper for the Intrinsic Circadian Clock"

**This PDF file includes:**

Materials and Methods
Figs. S1 to S8
Tables S1 to S2
Captions for Movies S1 to S4
References

**Other Supplementary Materials for this manuscript include the following:**

Movies S1 to S4

**Materials and Methods**

Mice

Source of animals was described in Table S1. All the mouse lines used in this work were backcrossed and maintained in C57BL/6J background. To generate experimental mouse model, *Nms-Ift88*$^{-/-}$, *Nms-Ift20*$^{-/-}$ or *Nms-Smo*$^{-/-}$ mice were generated by crossing *Nms-Cre* mice to *Ift88*$^{fl/fl}$, *Ift20*$^{fl/fl}$ or *Smo*$^{fl/fl}$ mice for at least two generations, respectively. The littermate *Nms-Cre* mice served as controls. *Nms-Ift88*$^{-/-}$ mice carrying Per2::Luc reporter were generated by crossing *Nms-Ift88*$^{-/-}$ to *Per2::Luc* mice. Likewise, *Rosa26-stop-tdTomato* mice were crossed with *Nms-Cre* or *Vip-Cre* mice for at least two generations to generate *Nms-tdTomato* or *Vip-tdTomato* mice, respectively. All animals were housed in specific-pathogen free facilities on a standard 12-h-light (50 lux)/12-h-dark cycle (LD, light on at 7:00 and light off at 19:00), with controlled temperature (22°C) and humidity (40%~60%). Regular food and water were provided *ad libitum*. Postnatal pups were obtained from natural matting and were maintained together with their mothers before being sacrificed at P2–P3. All animal experiments were performed with the approval of the Institutional Animal Care and Use Committee of Academy of Military Medical Sciences, Beijing, China (IACUC-DWZX-2020-508).

Locomotor activity recording

Adult female mice (8–10 weeks) were individually housed in cages equipped with a running wheel. Mice were initially entrained to LD cycle for at least 2 weeks before being exposed to constant darkness (DD) cycles or experimental jet-lag condition. Data was collected and analyzed using ClockLab (Actimetrics).

Intraventricular infusion of Vismodegib

Drug was continuously delivered to the SCN via osmotic pump. In brief, the tip of the infusion cannula (Brain Infusion Kit 2, Alzet) was directed stereotaxically to the SCN and the osmotic pump (pump model 1002, infused at 0.25 μl/h, Alzet) containing vehicle or 5 mM Vismodegib (Selleck) was placed in a subcutaneous pouch by surgery under anaesthesia. The vehicle solution used in administration was artificial cerebrospinal fluid (aCSF, 0.15 M NaCl, 3 mM KCl, 1.4 mM CaCl2, 0.8 mM MgCl2, 0.8 mM Na2HPO4, 0.2 mM NaH2PO4) containing 30% PEG300 and 5%

Tween 80. Following the surgery, mice were placed on a heating pad until weak up and were returned to their home cages. Three days after surgery, LD cycles were advanced by 8 h.

Light stimulation

Mice were entrained on a standard LD cycle for 2 weeks and were released to DD condition for 3 days. For light response experiment, the control, *Nms-Ift88*$^{-/-}$ and *Nms-Ift20*$^{-/-}$ mice were exposed to a 30-min light pulse (50 lux) at CT 22. Mice were sacrificed at CT 22 and CT 22.5 immediately under dim red light (n=3 for each time point), and the brains were collected followed by 4% paraformaldehyde fixative.

*Ex vivo* Culture of SCN slice

As reported previously (*36*), mice carrying Per2::Luc reporter were anesthetized and decapitated before onset of darkness, and their brains were rapidly removed. Coronal slices of the brain (300 µm) were made by Vibratome (VT1200S, Leica) in cold HBSS buffer (Gibco). SCN slices were then dissected into a 3 mm × 3 mm square with optic chiasm attached and cultured on top of the Millicell culture membrane (0.4 µm, Millipore) with 1.2 ml Dulbecco's modified Eagle's medium (DMEM, Gibco), supplemented with 2% B27 (Invitrogen), 1% penicillin/streptomycin (Gibco) and 0.1 mM Beetle Luciferin (Promega).

Bioluminescence recording and imaging

SCN cultures were maintained at 36°C in a lighttight incubator, and their bioluminescence was continuously monitored by LumiCycle luminometer (Actimetrics) equipped with photomultiplier tube (PMT) detector. For drug treatment, Vismodegib (5–20 µM, Selleck) was dissolved and added by changing pre-warmed fresh medium. For washout of Vismodegib, cultured slices were washed at least 3 times with pre-warmed medium and transferred to fresh medium containing luciferin. During the temperature entrainment assay, for normal body temperature cycles, SCN slices carrying Per2::Luc reporter were removed from LumiCylce luminometer and transferred to a homothermal incubator set at 36°C for 12 h, and then the cultures were moved to another incubator set at 38.5°C for 12 h; for oppositely phased temperature cycles, the cultures were transferred to the incubator set at 38.5°C for 12 h, and were then moved to 36°C for 12 h. Three days after temperature entrainment, the cultures were recorded in LumiCyle at 36°C.

Bioluminescence imaging was generally performed as described above with modifications. Identical culture conditions for SCN slices were used except that a higher concentration of luciferin (1 mM) was added for imaging. Culture dish was sealed and placed on the stage of an inverted microscope with 10× objective lens (Nikon Eclipse Ti-E) in a dark room. The stage temperature was set at 36°C throughout the experiment. A CCD camera (EA4710V-BV, Raptor, UK) operating at -80°C was mounted with the microscope for image capture. Images of 60 min exposure duration were collected continuously. 2 μM TTX or 10 μM Vismodegib was dissolved and added by changing pre-warmed fresh medium. TTX was washed out by moving the culture insert to a fresh medium for three times.

Visualization of cilia in living SCN neurons

To isolate single-cell SCN neurons, SCN slices were dissected from postnatal mice (P2–P3, n=30) as described above, and were enzymatically digested through Neural Tissue Dissociation Kit (130-092-628, Miltenyi Biotec) according to the manufacturer's instructions. In brief, dissected SCN tissues were transferred into the tube containing pre-heated enzyme mix and gently digested under gentleMACS Program 37C_NTDK_1 with the gentleMACS™ Dissociator (Miltenyi Biotec). After termination of the program, the samples were resuspended with PBS and then applied to a MACS SmartStrainer (70 μm, Miltenyi Biotec). Isolated viable neurons were then cultured with DMEM medium containing 2% B27 in a confocal dish which had been coated a layer of poly-lysine in advance for adherence. When the cells were completely adherent, cilia-targeted cADDis BacMam (D0211G, Montana Molecular) was transduced to label primary cilia in live cells according to manufacturer's recommendation (*37*). Time-lapse images were collected every 10 min for at least 48 hours, using a 60×/1.40 oil objective lens on an inverted fluorescence microscope (Nikon Eclipse Ti-E) with an Ultra View spinning-disc confocal scanner unit (PerkinElmer) at 37°C with 5% $CO_2$.

Immunostaining

Mice brains and peripheral tissues were fixed in 4% paraformaldehyde and embedded in OCT compound (Sakura) or paraffin. Paraffin and frozen sections were cut at 3 μm and 6 μm, respectively. Sections were blocked with 3% normal goat serum and 0.1% Triton X-100 in PBS for 1 h. After overnight incubation of primary antibodies at 4°C, sections were washed 3 times with PBS and stained with secondary antibodies at room temperature for 1h. DNA was stained with Hoechst 33342 (1:1000, H3570; Thermo Fisher Scientific). Sections were washed with PBS three times and mounted for confocal imaging. Immunofluorescence images were obtained using 63×/1.40 oil objective on Zeiss LSM 880 and analyzed by ZEN2.1 SP2 Black version 13.0.2.518 (ZEISS). To stain AVP neurons, mice were pretreated with intracerebroventricular injections of colchicine (20 µg in 1 µl saline), and 48 hours later, mice were sacrificed at approximately ZT 0. For NeuN staining, paraffin slides were dewaxed with xylol and rehydrated through a graded alcohol series. The endogenous peroxidase was inactivated by immersing in a 3% hydrogen peroxide solution for 10 min. Antigen retrieval was carried out by boiling the slides in 10 mM sodium citrate buffer (pH 6.0) for 10 min. The slides were blocked with 10% normal goat serum for 10 min at room temperature and were then incubated with anti-NeuN antibody at 37°C for 1 h. Immunoreactivities were visualized by HRP DAB Detection kit (ZLI-9018, ZSGB-BIO). Then, slides were stained with hematoxylin for 2 min to stain nucleus. All steps were followed by appropriate washing in PBS. Finally, slides were mounted with composite resin and images were captured using NanoZoomer Digital Pathology system (Hamamatsu Photonics).

Antibodies

Antibodies used in this work are as follows: rabbit anti-A cyclase III (AC III, 1:400, sc-588; Santa Cruz Biotechnology), rabbit anti-mCherry (1:400, ab167453; Abcam), rabbit anti-ARL13B (1:400, 17711-1-AP; Proteintech), rabbit anti-Cry1 (1:100, A302-614A; Bethyl), mouse anti-AVP (1:200, sc-390702; Santa Cruz Biotechnology), Goat anti-Rabbit Alexa Fluor 488 (1:100, A11034; Thermo Fisher Scientific), Goat anti-Rabbit Alexa Fluor 546 (1:100, A11035; Thermo Fisher Scientific), Goat anti-Mouse Alexa Fluor 546 (1:100, A11030; Thermo Fisher Scientific), rabbit anti-NeuN (1:500, ab12452; Abcam), rabbit anti-SMO (1:1000, A3274; ABclonal), rabbit anti-IFT88 (1:500, 13967-1-AP; Proteintech), rabbit anti-IFT20 (1:1000, 13615-1-AP; Proteintech), rabbit anti-α-tubulin (1:5000, PM054; MBL).

SCN tissue clearing and two-photon microscopy imaging

SCN slices were dissected as described above after perfusion and processed tissue clearing using SHIELD method (*38*). Passive immunostaining of primary cilia with AC III in the clearing SCN slices was applied according to the procedure from Nuohai Life Science, China. Fluorescent images of the whole SCN were obtained using two-photon microscopy (Olympus, FV1200MPE) with 20× objective lens. The 3D structures of the SCN were reconstructed by using the Z-stack function of Imaris software (Imaris 9.2.4, Bitplane).

RNA extraction and quantitative real-time PCR (qRT-PCR)

Total RNA of SCN tissues (each sample was pooled from 6 mice) was extracted using TRIzol reagent (93289; Sigma-Aldrich) and was reverse-transcribed to complementary DNA (cDNA) using the PrimeScript™ RT Master Mix (RR036A, Takara). cDNA was used to determine relative mRNA levels by qRT-PCR running on a Roche StepOnePlus Real-Time PCR System, and *Gapdh* (housekeeping gene) served as control. Data were analyzed using StepOnePlus software. The primer sequences of targeted genes are listed in Table S2.

RNAscope in situ hybridization

RNAscope experiment was carried out using RNAscope Multiplex Fluorescent Reagent Kit (323100, ACDbio) in accordance with manufacturer's instructions. The target probes of *Cry1* (500031-C3, ACDbio), *Per1* (438751, ACDbio), *cFos* (316921-C3, ACDbio), *Shh* (314361-C2, ACDbio), *Ptch1* (402811-C2, ACDbio) and *Gli1* (311001-C2, ACDbio) were used to detect mRNA expression, respectively.

Nissl staining

Paraffin sections were dewaxed by xylol and graded ethanol, and then dipped in 0.2% cresyl violet (Solarbio) solution for about 5 min to stain nissl bodies. After rinsing in distilled water, slides were differentiated in 75% and 90% ethanol for 2 min each. Then, the slides were dehydrated in absolute ethanol for 3 min and were dipped in the 100% xylol for 5 min. Finally, slides were mounted with composite resin and images were captured using NanoZoomer Digital Pathology system (Hamamatsu Photonics).

Quantification and statistical analysis

All the data were expressed as mean ± SEM. The diurnal data was employed to fit the data into a smooth curve in GraphPad Prism8. The data was considered diurnal oscillation by Jonckheere-Terpstra-Kendall (JTK) analysis with a $P$-value of less than 0.05.

Locomotor activity was analyzed by ClockLab (Actimetrics). Behavioral period was calculated by Chi-squared periodogram. Phase shifts were quantified as time differences between regression lines of activity onsets or offsets before and after the light application. To calculate 50% phase shift values (PS50), sigmoidal curve, $Y = \text{Bottom} + (\text{Top} - \text{Bottom})/(1 + 10^{(\log \text{PS50} - X)\ \text{HillSlope}})$, was fitted to onset or offset values, using the interpolate function in GraphPad Prism8 (*39*).

Bioluminescence recording was analyzed by using Lumicycle Analysis program (Actimetrics). The first cycle was excluded from raw data due to high transient luminescence upon medium change. Period and amplitude of the wave were accurately calculated by using the FFT-NLLS function of the online BioDare2 suite (https://biodare2.ed.ac.uk) (*40*).

Single-cell bioluminescence analysis was performed in a similar manner as previously described (*19, 28*). 16-bit time-series images were imported and analyzed in Volocity 6.0 (Perkin Elmer). The cosmic ray noise was removed using median filter, and the bioluminescence value of single cell in time-series images were outlined individually within a region of interest (ROI, 10×10 pixel). ROI data was exported into Excel (Microsoft) for background subtraction and were detrended using Hodrick-Prescott filter with parameter $\lambda = 16588.8$ in EViews 10 (IHC Markit). Then the detrended data was smoothed using an adjacent averaging method within 2 hours, and was curve fitted to a sine wave with exponentially decaying amplitude in Igor Pro (WaveMetrics):

$$L(t) = A_0 \cdot e^{-t/k} \cdot \sin(2\pi t/\tau - \varphi)$$

where $L$ is luminescence intensity, $t$ is time, $A_0$ is amplitude at $t = 0$, $k$ is time constant for exponential decay of amplitude, $\tau$ is circadian period, and $\varphi$ is circadian phase. The fitted data was imported into OriginPro 2021 (OriginLab) to plot the trajectory of each cell bioluminescence.

For heatmaps plotting in Fig. 3C and 4D, fitted ROI data was calculated for z-score normalization and plotted in GraphPad Prism8 according to their phase orders (earlier phases were placed at the top). Circadian phase of each cell during the third day (midpoint) of continuous recording was estimated in BioDare2, and the phase clustering was assessed by Rayleigh's test and plotted in Oriana4 (Kovach Computing Services, Anglesey, UK).

For movie presentation, filtered image sequence was processed with contrast enhancement in Volocity 6.0 (Perkin Elmer), and was exported to Quicktime (Apple) for conversion to H264 videos.

Statistical significance was calculated with SPSS. We used two-tailed Student's *t*-test to compare two groups of normally distributed data. One or two-way analysis of variance (ANOVA) was used to compare multiple groups of normally distributed data followed by Bonferroni's or Dunnett's post hoc comparisons. Kruskal-Wallis test was performed to compare multiple groups of data that do not meet normal distribution. Watson-Wheeler test was performed to analyze the uniformity of phase distribution of Rayleigh plots. The test methods, the significance values and the number of experimental replicates for each experimental group were noted in the figure legends. The various asterisk abbreviations denote *: $p < 0.05$, **: $p < 0.01$, ***: $p < 0.001$, ns: not significant.

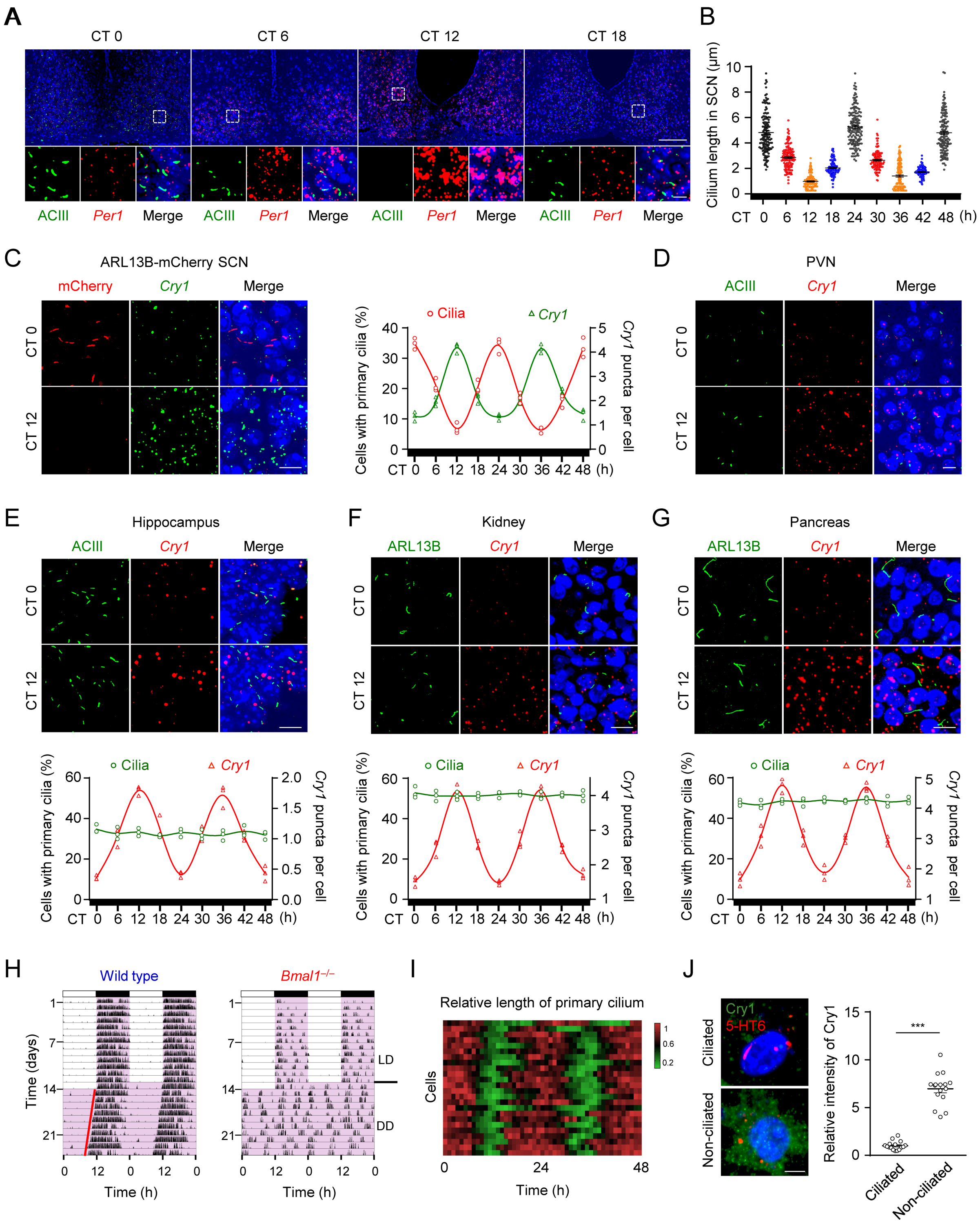


**Fig. S1. Primary cilia specifically exhibit circadian rhythmic growth patterns in the SCN.**

(**A**) Representative images of primary cilia and expression of clock gene *Per1* in the SCN during constant darkness (DD) cycle in Fig. 1F. SCN coronal sections were stained with anti-ACIII antibody (green), *Per1* RNAscope probes (red) and Hoechst (blue). Insets show zoomed-in views of the boxed regions in the core of the SCN. Scale bars, 100 μm (main image) and 10 μm (magnified region). (**B**) Quantitative analysis of the cilium length in (A). Each dot represents one cell. (**C**) The percentage of cells with primary cilia or *Cry1* RNAscope signals in the SCN of transgenic mouse expressing ARL13B-mCherry fusion protein were quantified during DD cycle ($n$ = 3 mice per time point). SCN coronal sections were stained with anti-mCherry antibody (red), *Cry1* RNAscope probes (green) and Hoechst (blue). Representative images at CT 0 and CT 12 are shown. Scale bar, 10 μm. (**D**) Representative images of primary cilia and expression of clock gene *Cry1* at CT 0 and CT 12 in Fig. 1G. Scale bar, 10 μm. (**E** to **G**) The percentage of cells with primary cilia or *Cry1* RNAscope signals in the hippocampus (E), kidney (F) and pancreas (G) were quantified during DD cycle (n = 3 mice per time point). The sections were stained with ciliary marker (ACIII or ARL13B, green), *Cry1* RNAscope probes (red) and Hoechst (blue). Representative images at CT 0 and CT 12 are shown. Scale bars, 10 μm. (**H**) Representative double-plotted actograms of wheel-running activities in wild type and $Bmal1^{-/-}$ mice. (**I**) Representative heatmap for the rhythmicity of primary cilium in Fig. 1I. Each row represents individual neuron. The length of primary cilia was normalized to the maximum length. (**J**) Left, representative images of ciliated or non-ciliated SCN neurons in cultured medium. The SCN neurons isolated from postnatal mice were transduced with modified baculovirus encoding a mCherry 5-HT6, and then were co-stained with anti-Cry1 antibody (green) and Hoechst (blue). Scale bar, 5 μm. Right, quantification of the relative intensity of Cry1 in isolated ciliated or non-ciliated SCN neurons ($n$ = 16). All data are presented as mean ± SEM. Statistics indicate significance by unpaired $t$-test (J). ***$P < 0.001$.

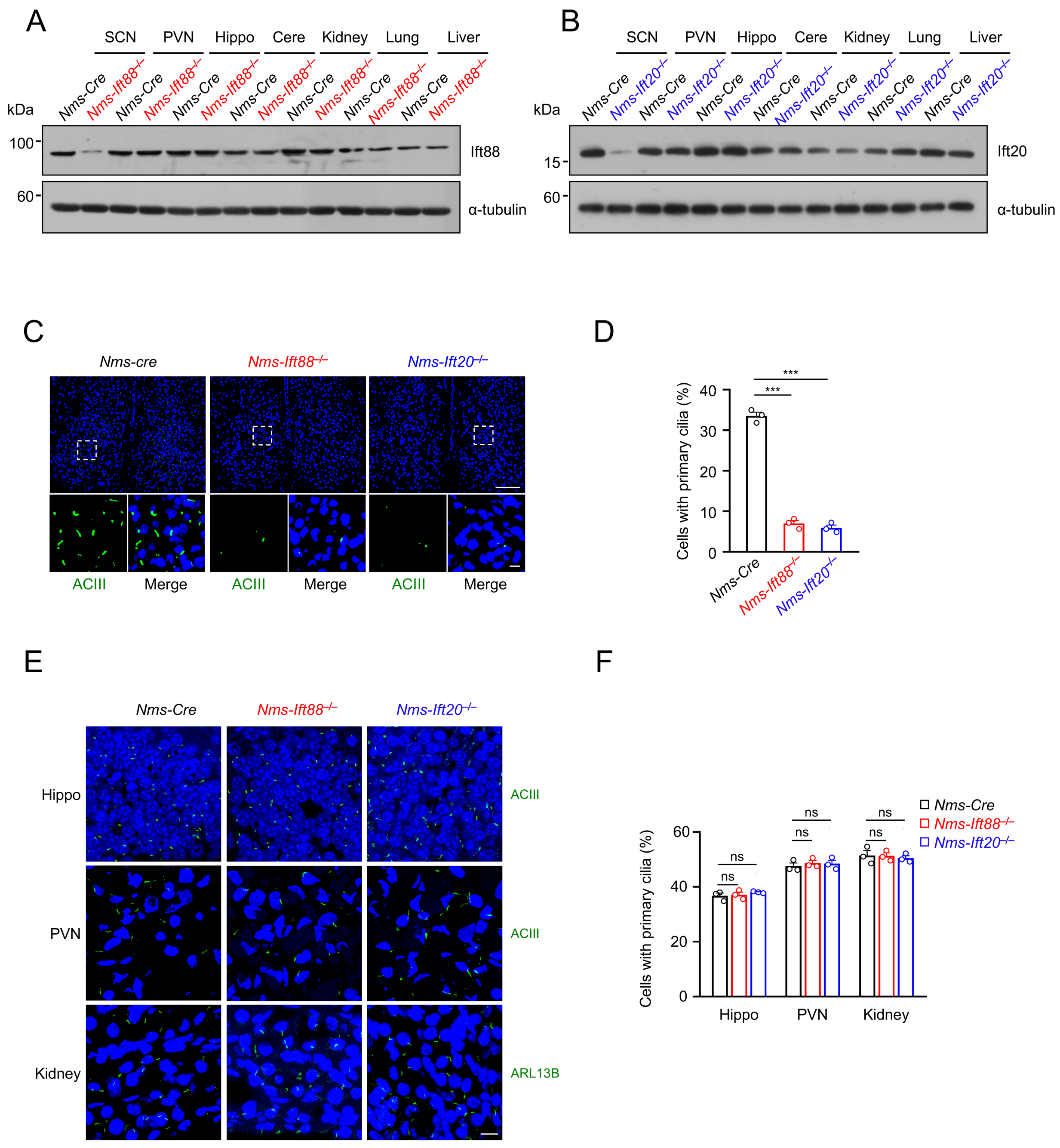


**Fig. S2. Primary cilia in the SCN are abolished in *Nms-Ift88*$^{-/-}$ or *Nms-Ift20*$^{-/-}$ mice.**

(**A** and **B**) Immunoblot analysis of protein extracts from *Nms-Cre*, *Nms-Ift88*$^{-/-}$ or *Nms-Ift20*$^{-/-}$ mice by using indicating antibodies. α-tubulin was used as a loading control. PVN, paraventricular nucleus. Hippo, hippocampus. Cere, cerebellum. (**C**) Representative images of primary cilia in the

SCN from *Nms-Cre*, *Nms-Ift88*$^{-/-}$ or *Nms-Ift20*$^{-/-}$ mice at ZT 0. SCN coronal sections were stained with anti-ACIII antibody (green) and Hoechst (blue). Insets show zoomed-in views of the boxed regions in the core of the SCN. Scale bars, 100 μm (main image) and 10 μm (magnified region). (**D**) Quantitative analysis of the ratio of ciliated cells in (C) ($n$ = 3). (**E**) Representative images of cilia in the hippocampus, PVN and kidney from *Nms-Cre*, *Nms-Ift88*$^{-/-}$ or *Nms-Ift20*$^{-/-}$ mice at ZT 0. The sections were stained with anti-ACIII or anti-ARL13B antibody (green) and Hoechst (blue). Scale bar, 10 μm. (**F**) The percentage of cells with primary cilia in (E) ($n$ = 3). All data are presented as mean ± SEM. Statistics indicate significance by one-way ANOVA with Dunnett correction (D), two-way ANOVA with Dunnett correction (F). ***$P < 0.001$, ns, not significant.

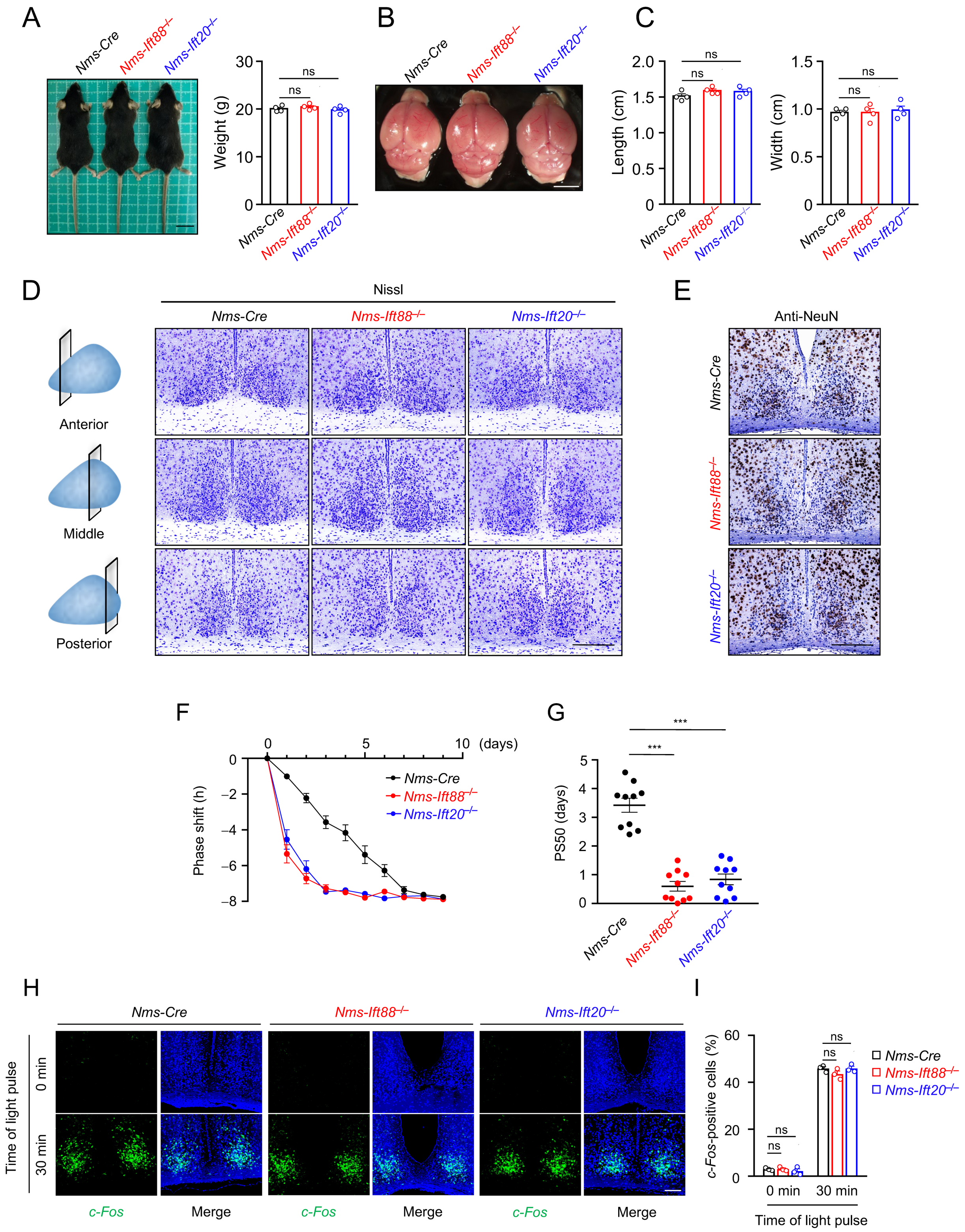


**Fig. S3. The development, locomotor activity and light response analysis of SCN$^{cilia\text{-}null}$ mice.**

(**A**) Representative picture and weight analysis for *Nms-Cre* and $SCN^{cilia\text{-}null}$ mice at 10-week old (*n* =4). Scale bar, 2 cm. (**B**) Representative picture of the brain for *Nms-Cre* and $SCN^{cilia\text{-}null}$ mice. Scale bar, 4 mm. (**C**) Quantitative analysis of length and width of the brains in (B) (*n* = 4). (**D**) SCN structure analyses of *Nms-Cre* and $SCN^{cilia\text{-}null}$ mice by Nissl staining. Scale bar, 200 µm. (**E**) Immunohistochemistry staining of the SCN from *Nms-Cre* and $SCN^{cilia\text{-}null}$ mice for NeuN (neuronal marker). Scale bar, 200 µm. (**F**) Line graphs showing the daily phase shift of wheel-running activities after an 8 h delay in Fig. 2F. (**G**) PS50 values after an 8 h delay in Fig. 2F. (**H**) Representative images of *c-Fos* RNAscope fluorescence in the SCN from *Nms-Cre* and $SCN^{cilia\text{-}null}$ mice. Animals were exposed to 30 min light pulse at CT 22 and sacrificed. SCN coronal sections were stained with *c-Fos* RNAscope probes (green) and Hoechst (blue). Scale bar, 100 µm. (**I**) Quantification of the percentage of *c-Fos* positive cells in (H) (*n* = 3). All data are presented as mean ± SEM. Statistics indicate significance by one-way ANOVA with Dunnett correction [(A), (C) and (G)], two-way ANOVA with Dunnett correction (I). ***$P < 0.001$, ns, not significant.

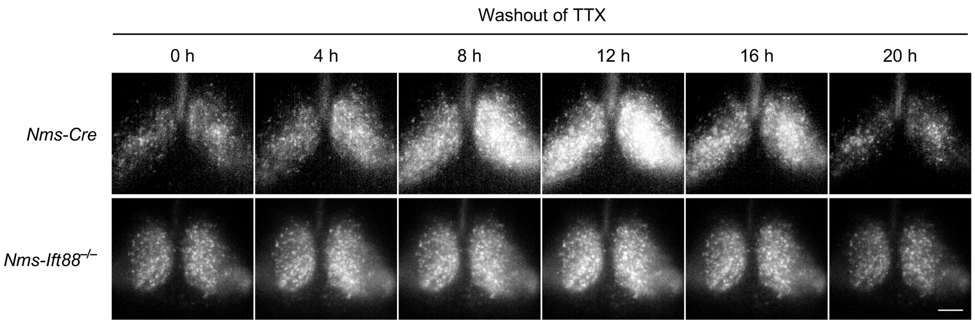


**Fig. S4. Phase order in *Nms-Ift88*$^{-/-}$ SCN fails to recover after the washout of TTX.**

Representative time-lapse images of the SCN slice from *Nms-Cre* and *Nms-Ift88*$^{-/-}$ mice after TTX washout in Fig. 3, A and B. The trough time of SCN bioluminescence on day 2 was set to time 0. Scale bar, 200 μm.

A

B

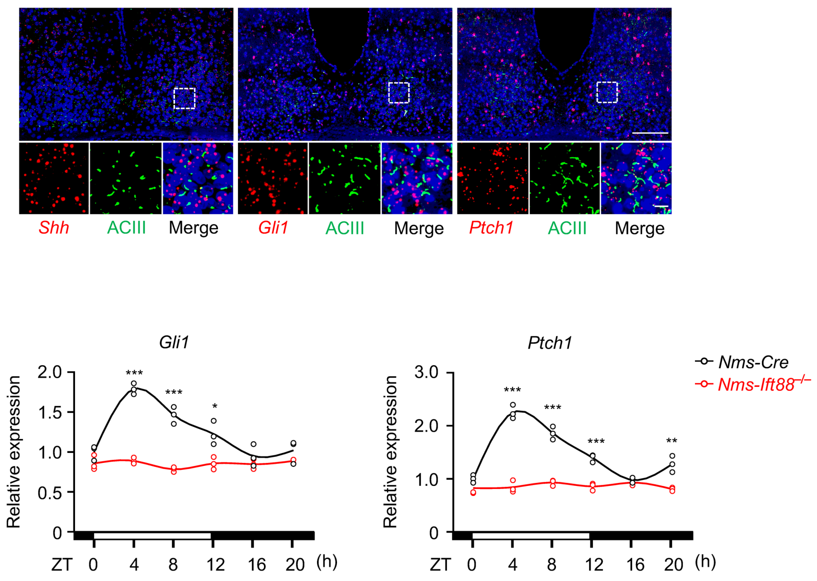


**Fig. S5. The rhythmicity of Hedgehog signaling is abolished in *Nms-Ift88*$^{-/-}$ mice.**

(**A**) Representative images of primary cilia and expression of *Shh*, *Gli1* and *Ptch1* in the SCN. SCN coronal sections were stained with anti-ACIII antibody (green) and *Shh*, *Gli1*, *Ptch1* RNAscope probes (red) and Hoechst (blue). Insets show zoomed-in views of the boxed regions in the core of the SCN. Scale bars, 100 μm (main image) and 10 μm (magnified region). (**B**) Quantitative real-time PCR analysis of Hedgehog signaling target genes *Gli1* and *Ptch1* in the SCN from *Nms-Cre* and *Nms-Ift88*$^{-/-}$ mice. SCN were collected at 4 h intervals across the LD cycle ($n$ = 3 independent experiments). Data are presented as mean ± SEM. Statistics indicate significance by unpaired *t*-test. *$P < 0.05$, **$P < 0.01$, ***$P < 0.001$.

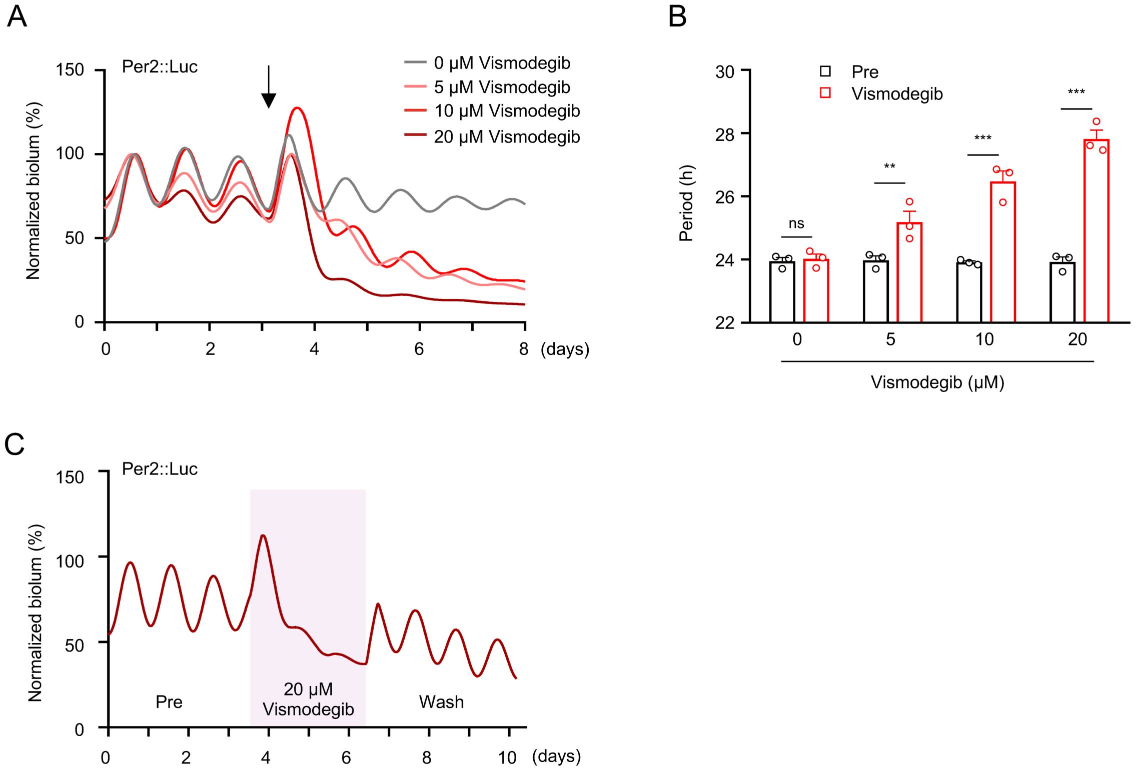


**Fig. S6. Inhibition of Hedgehog signaling with Vismodegib disrupts Per2::Luc oscillations.**

(**A**) Representative records of Per2 bioluminescence in Per2::Luc SCN slices treated with Vismodegib. Bioluminescence was normalized to the first peak. Arrow indicates Vismodegib treatment. (**B**) Quantitative analysis of the period of Per2 bioluminescence in (A) ($n = 3$). Data are presented as mean ± SEM. Statistics indicate significance by two-way ANOVA with Bonferroni correction. $^{**}P < 0.01$, $^{***}P < 0.001$, ns, not significant. (**C**) Representative records of Per2 bioluminescence in Per2::Luc SCN slices treated with Vismodegib, followed by washout. Bioluminescence was normalized to the first peak.

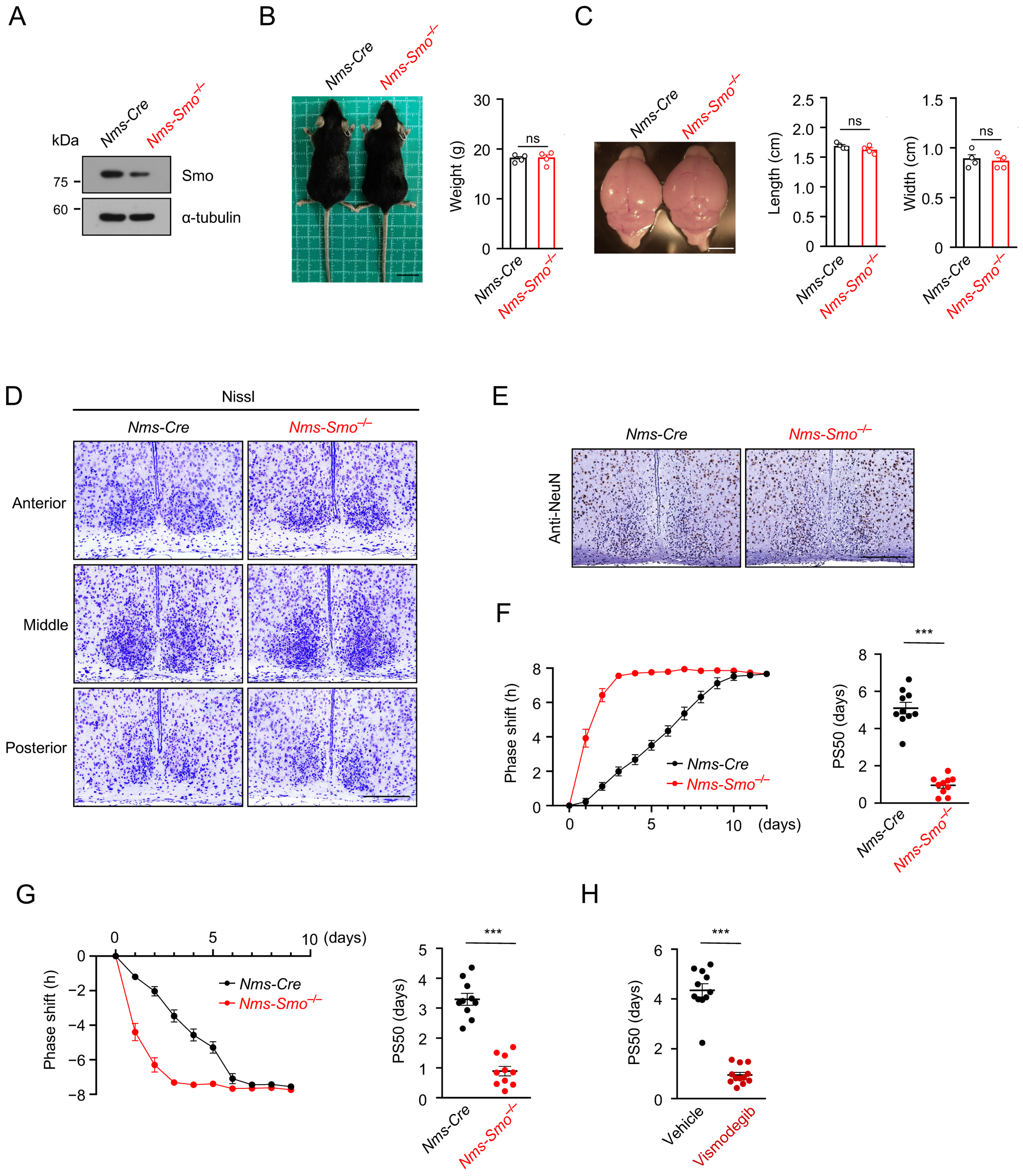


**Fig. S7. The development and locomotor activity analysis of *Nms-Smo*$^{-/-}$ mice.**

(**A**) Immunoblot analysis of SCN protein extracts from *Nms-Cre* and *Nms-Smo*$^{-/-}$ mice by using indicating antibodies. α-tubulin was used as a loading control. (**B**) Representative picture and weight analysis for *Nms-Cre* and *Nms-Smo*$^{-/-}$ mice at 8-week old ($n$ = 4). Scale bar, 2 cm. (**C**) Representative picture and quantitative analysis of length and width of the brains from *Nms-Cre* and *Nms-Smo*$^{-/-}$ mice ($n$ = 4). Scale bar, 4 mm. (**D**) SCN structure analyses of *Nms-Cre* and *Nms-Smo*$^{-/-}$ mice by Nissl staining. Scale bar, 200 μm. (**E**) Immunohistochemistry staining of the SCN from *Nms-Cre* and *Nms-Smo*$^{-/-}$ mice for NeuN. Scale bar, 200 μm. (**F**) Line graphs and PS50 values showing the daily phase shift of wheel-running activities after an 8 h advance in Fig. 4A. (**G**) Line graphs and PS50 values showing the daily phase shift of wheel-running activities after an 8 h delay in Fig. 4A. (**H**) PS50 values after an 8 h advance in Fig. 4B. All data are presented as mean ± SEM. Statistics indicate significance by unpaired *t*-test [(B), (C), (F), (G) and (H)]. ***$P$ < 0.001, ns, not significant.

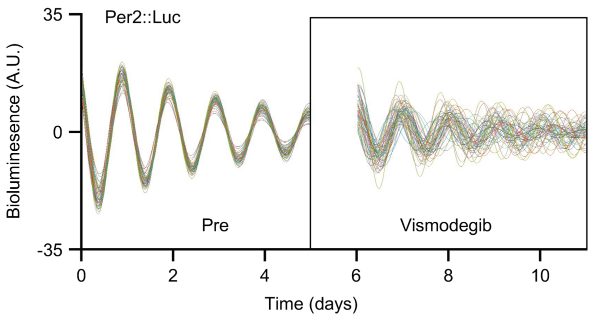


**Fig. S8. Single-cell bioluminescence analysis of SCN slices with Vismodegib treatment.**

Representative records of single-cell Per2 bioluminescence in Fig. 4D.

**Table S1** List of Mouse Source. Related to Materials and Methods.

| Genotype | Stock No. | Source |
|---|---|---|
| *Ift88* $^{fl/fl}$ | 022409 | Jackson Laboratory |
| *Ift20* $^{fl/fl}$ | 012565 | Jackson Laboratory |
| *Smo* $^{fl/fl}$ | 004526 | Jackson Laboratory |
| *Nms-Cre* | 027205 | Jackson Laboratory |
| *Vip-Cre* | 031628 | Jackson Laboratory |
| *ARL13B-mCherry* | 027967 | Jackson Laboratory |
| *Rosa26-stop-tdTomato* | T002249 | Gempharmatech |

**Table S2** Quantitative PCR Primer Sequences. Related to Materials and Methods.

| Gene | Fw | Rv |
|---|---|---|
| *Bmal1* | GCAGTGCCACTGACTACCAAGA | TCCTGGACATTGCATTGCAT |
| *Clock* | ACCACAGCAACAGCAACAAC | GGCTGCTGAACTGAAGGAAG |
| *Per1* | ACCAGCGTGTCATGATGACATA | GTGCACAGCACCCAGTTCCC |
| *Cry1* | CAGACTCACTCACTCAAGCAAGG | TCAGTTACTGCTCTGCCGCTGGAC |
| *Gli1* | CCAAGCCAACTTTATGTCAGGG | AGCCCGCTTCTTTGTTAATTTGA |
| *Ptch1* | AAAGAACTGCGGCAAGTTTTTG | CTTCTCCTATCTTCTGACGGGT |
| *Gapdh* | AGGTCGGTGTGAACGGATTTG | TGTAGACCATGTAGTTGAGGTCA |

**Movie S1.**

3D assessment of primary cilia in the SCN. Related to Fig. 1B.

**Movie S2.**

The primary cilium in the SCN neuron exhibits circadian rhythmic oscillation. Related to Fig. 1I.

**Movie S3.**

Bioluminescence imaging of *Nms-cre* SCN slice after the washout of TTX. Related to Fig. 3A.

**Movie S4.**

Bioluminescence imaging of *Nms-Ift88*$^{-/-}$ SCN slice after the washout of TTX. Related to Fig. 3B.
